# Correction of susceptibility distortion in EPI: a semi-supervised approach with deep learning

**DOI:** 10.1101/2022.07.12.499591

**Authors:** Antoine Legouhy, Mark Graham, Michele Guerreri, Whitney Stee, Thomas Villemonteix, Philippe Peigneux, Hui Zhang

## Abstract

Echo planar imaging (EPI) is the most common approach for acquiring diffusion and functional MRI data due to its high temporal resolution. However, this comes at the cost of higher sensitivity to susceptibility-induced *B*_0_ field inhomogeneities around air/tissue interfaces. This leads to severe geometric distortions along the phase encoding direction (PED). To correct this distortion, the standard approach involves an analogous acquisition using an opposite PED leading to images with inverted distortions and then non-linear image registration, with a transformation model constrained along the PED, to estimate the voxelwise shift that undistorts the image pair and generates a distortion-free image. With conventional image registration approaches, this type of processing is computationally intensive. Recent advances in unsupervised deep learning-based approaches to image registration have been proposed to drastically reduce the computational cost of this task. However, they rely on maximizing an intensity-based similarity measure, known to be suboptimal surrogate measures of image alignment. To address this limitation, we propose a semi-supervised deep learning algorithm that directly leverages ground truth spatial transformations during training. Simulated and real data experiments demonstrate improvement to distortion field recovery compared to the unsupervised approach, improvement image similarity compared to supervised approach and precision similar to TOPUP but with much faster processing.

## 1 Introduction

Echo planar imaging (EPI) is the most common approach for acquiring diffusion and functional MRI data due to its high temporal resolution which both reduces the influence of motion and allows the acquisition of a large number of volumes in a time frame amenable to neuroscientific and clinical research. This is, however, at the cost of higher sensitivity to susceptibility-induced *B*_0_ field inhomogeneities around interfaces of air, bone, and soft tissue. This leads to severe geometric distortions in the form of local expansions or contractions along the phase encoding direction (PED), breaking alignment with the corresponding anatomical scans.

To tackle this problem, the strategy that has proved most effective is to acquire an extra scan with identical settings, except for an opposite PED [1]. It produces an analogous image with reverse distortion: expansions where there were contractions and vice versa. One can then apply non-linear image registration, with a transformation model constrained along the PED, to estimate the shift in voxel coordinates that undistorts the image pair and generates a distortion-free image.

Standard implementations of this strategy, e.g. TOPUP [3] in FSL, are computationally intensive. They rely on traditional image registration methods that align each new pair of images with a separate iterative optimization. A comparison of such algorithms is available in [2]. Recently, image registration based on deep learning (DL) architectures, convolutional neural networks (CNN) in particular, have been developed. With the investment of an upfront cost during training, test images can be registered in one-shot almost instantaneously. This has led to the development of fast EPI distortion correction with DL-powered image registration [4]. Based on VoxelMorph [12], an unsupervised framework, this approach, similar to traditional registration, relies on maximizing an intensity similarity measure, known to be suboptimal surrogate measures of image alignment [5]. Also, it does not integrate intensity modulation to account for signal stretchings and pile-ups associated to geometric expansions and contractions.

To address those limitations, we propose a semi-supervised DL algorithm that directly leverages ground truth spatial transformations during training. We hypothesise that constraining DL model training with the most direct representations of spatial correspondence will significantly improve the fidelity of the recovered spatial correspondence during testing. Also we integrate Jacobian intensity modulation when constructing the undistorted images.

We performed two experiments to evaluate the proposed method against the unsupervised approach similar to [4], but also against a fully transformation supervised one. The first experiment makes use of real data to assess the practical performance. For this experiment, transformations produced by TOPUP are used as ground truth for training and testing. The second experiment makes use of simulated data, generated by DW-POSSUM [1], a realistic Spin-Echo EPI simulator. This sets up a scenario where we have genuine ground truth for both undistorted image and the distortion-inducing deformation, and allows the comparison with TOPUP.

## 2 Background

### 2.1 Distortion model

Susceptibility-induced EPI geometric distortion is well understood [7]. For such multi-slice acquisitions, distortion due to *B*_0_ field inhomogeneities is negligible along the frequency encoding direction. Its effect can thus be parametrized by a unidirectional deformation field *V* shifting voxel coordinates *x* along the PED. For the typical PED from posterior to anterior, we have *T*_+_ (*x*) = *x* + *V*(*x*). If the PED is reversed, an opposite displacement field will result: *T*_−_(*x*) = *x* – *V*(*x*). Henceforth, we will refer to this as the opposite symmetry constraint for the two displacement fields.

Without loss of generality, we will refer to *T*_+_ as the forward transform, and the corresponding distorted image as *I*_+_. Likewise, *T*_−_ will be referred to as the backward transform and the corresponding distorted image as *I*_−_. The latent undistorted image will be denoted as *I*. Following [8], it can be expressed both in terms of the forward and backward images following:

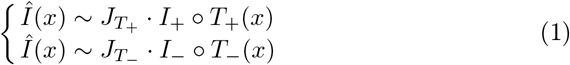

where *J*_*T*_+__ (resp. *J*_*T*_−__) is the Jacobian determinant of *T*_+_ (resp. *T*_−_), ◦ denoting composition and · element-wise multiplication. *J_T_* encodes the local expansion (if |*J_T_*| ∈ [1, +∞)) or contraction (if |*J_T_*| ∈ (0,1]) properties of the transformation and will modulate the resulting intensities accordingly [9].

### 2.2 Distortion correction using image registration

Under the above distortion model, it is evident image registration can be used to estimate the transformations for correcting the distorted image pair. Eq. (1) suggests the correction can be formulated as the following image registration problem:

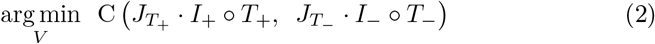

where C is a dissimilarity criterion between the two corrected images (from *I*_+_ and *I*_−_ respectively). The mean squared error (MSE) between intensities is particularly well suited in this case since we are dealing with the same subject, the same modality, the same acquisition parameters (with the exception of PED); we therefore expect almost identical intensities at endpoint. The sought transformations (*T*_+_ and *T*_−_) are parametrized by *V* which, even though constrained to be unidirectional, can have a large number of degrees of freedom: up to the number of voxels of the image.

As noted in the introduction, this image registration problem is currently typically solved via computationally intensive iterative optimization, with each new image pair solved completely independently. Recent advances in DL-based image registration have recently been leveraged to substantially accelerate this task by replacing iterative optimization with a one-shot computation [10] [11]. The idea behind DL-based image registration is to learn a model that can predict from an image pair the transformation that put them into correspondence. During training, model parameters are tuned to output an optimal transformation, for each training sample, that maximises either some similarity between transformed images (unsupervised) or the resemblance to the corresponding ground truth transformation (supervised). While the training may be computationally expensive, once completed, new image pairs can be registered almost instantaneously. The most popular publicly available implementation is VoxelMorph [12][13] which provides an unsupervised framework, making use of a U-Net convolutional neural network (CNN) architecture. This framework was recently exploited for EPI distortion correction [4]. However it has now been limited to optimisation over purely intensity-based losses and does not embed intensity modulation.

## 3 Method

We implement the proposed semi-supervised approach using the Voxelmorph framework. The framework must be adapted 1) to predict a spatial transform pair with opposite symmetry, 2) to constrain predicted spatial transforms to be unidirectional along the PED, 3) to support the image registration formulation represented by Eq. (2) that includes Jacobian intensity modulation, 4) to enable semi-supervision with ground truth spatial transforms, and 5) to handle weight maps that modulate the contribution of each voxels according to anatomical regions of interest. Points 3, 4 and 5 are not present in the work from [4]. The details of the model are described below.

### 3.1 Model architecture

The model is organised into three sequential blocks (Fig. 1). The first block takes a distorted image pair as input and outputs a unidirectional vector field which is used to produce a forward and a backward transformation. Those, together with the associated input images, are fed to a resampling block that reconstruct undistorted images by interpolating from transformed coordinates.

**Fig. 1.**
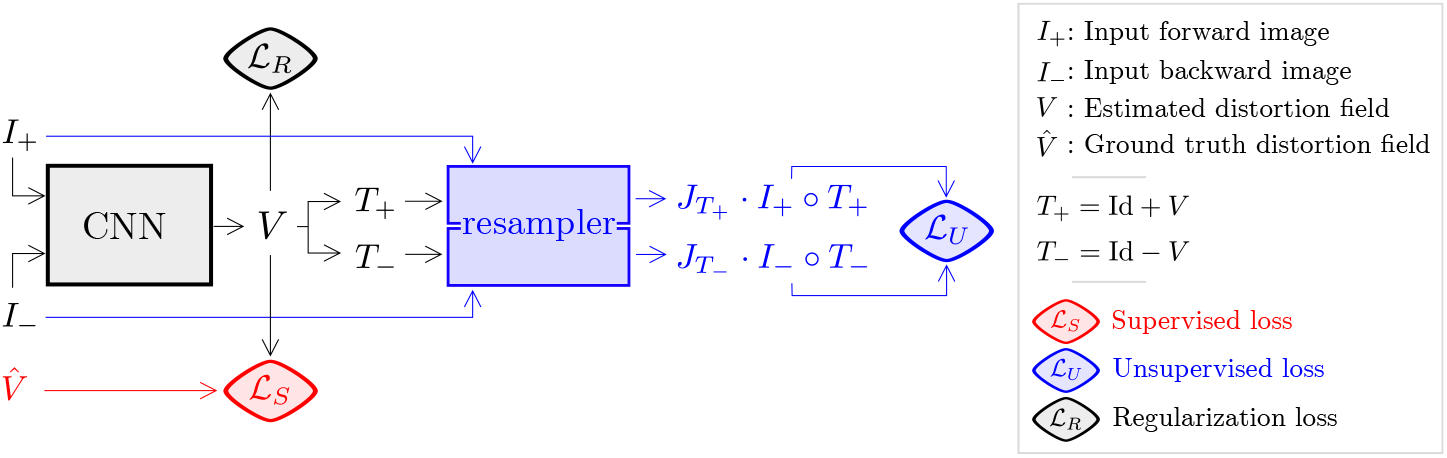
Diagram of the the proposed semi-supervised distortion correction network. Portions specific to unsupervised are highlighted in blue, whereas the ones associated to supervised are in red.

The CNN block is a U-Net as in VoxelMorph, but its input and output are different. Instead of any arbitrary image pair, it must be given a pair of analogously distorted images {*I*_+_,*I*_−_} as input. Instead of a vector field, it outputs a scalar field *V* characterizing constrained displacements along a single direction (PED). As in VoxelMorph, all the trainable parameters of the model are contained in this block.

To preserve the opposite symmetry constraint induced by the PED reversal, the pair of forward and backward transformations {*T*_+_,*T*_−_} are built from the estimated field *V* following: *T*_+_ = Id + *V* and *T*_−_ = Id – *V*.

Resampling block implements image warping. Unlike VoxelMorph and [4], our implementation includes intensity modulation, which allows us to account for signal pile-up in the presence of contraction and signal reduction in the presence of expansion. It takes *I*_+_ (resp. *I*_−_) and *T*_+_ (resp. *T*_−_) as input to produce undistorted, intensity-modulated images *J*_*T*_+__ · *I*_+_ ◦ *T*_+_ (resp. *J*_*T*_−__ · *I*_−_ ◦ *T*_−_). This requires computing the Jacobian determinant of the transformations.

### 3.2 Models

The model architecture described above is used to implement an unsupervised and a transformation supervised model, as baselines for comparison, and the proposed semi-supervised model.

A diagram illustrating the different models can be found in Fig. 1. The parts exclusive to the unsupervised model, comparable to [4], are highlighted in blue. It includes the resampling block that unwarps the input images with the estimated field allowing their comparison (unsupervised loss). The red parts are exclusive to the supervised model. It includes the ground truth field that is compared to the estimated one (supervised loss). The black parts are common to both. It includes the distortion field estimation (a regularization loss is computed on it). The semi-supervised model encompasses both the unsupervised and supervised components.

The unsupervised loss 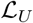 is an image similarity metric between the two undistorted images that have undergone Jacobian intensity modulation. As mentioned in Section 2.2 the MSE is well indicated and Eqn. 1 leads to:

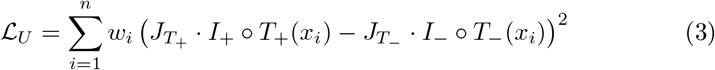

This loss is only required for the unsupervised and semi-supervised models.

The supervised loss is a distance between the estimated distortion field *V* and the ground truth one 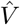. The MSE is also well adapted when dealing with displacement vectors as it corresponds to an average of geometrical distances, it’s a direct quantitative measure of the goodness of the registration:

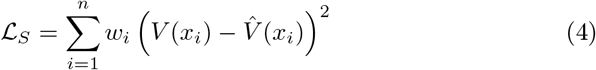

This loss is only required for the supervised and the semi-supervised model.

A regularization loss 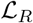 is also use on the estimated distortion field *V* to encourage smoothness:

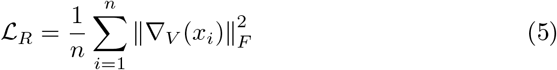

where ∇_*V*_ is the Jacobian of the field *V* and ||.||_*F*_ is the Frobenius norm. This loss is computed for all models.

For each models, the overall loss is a sum of the ones above, weighted to account for large order-of-magnitude differences in different loss terms.

To prevent the learning process to be influenced by meaningless information from background (which represent the majority of the image!), another kind of weighting, spatially this time, occurs when computing the unsupervised and the supervised losses. It corresponds to the *w_i_* in Eq. 3 and Eq. 4. The contribution of each voxels is modulated such as only areas of interest (typically just the brain) contribute to the loss, ignoring the background. This is not present in Voxelmorph and [4].

## 4 Evaluations

The idea is to evaluate how well, having processed a portion of a dataset with a regular tool (here TOPUP), one can use those to train a DL model that will rapidly process the rest of the dataset. Corrections from three DL registration approaches are engaged for comparison: unsupervised, transformation supervised and semi-supervised. We will perform experiments on two datasets: 1- A real dataset with TOPUP outputs as ground truths, 2- A synthetic dataset with absolute ground truths allowing to integrate TOPUP to the comparison.

### 4.1 Datasets

We acquired anatomical and diffusion weighted data as part of an ongoing study investigating memory learning and consolidation. We have, for 60 healthy subjects, T1-weighted images and distorted EPI b=0 image pairs from opposite PED acquisition that are antero-posterior (AP) and postero-anterior (PA).

Distorted EPI images have been processed through TOPUP to obtain undistorted images and associated distortion fields.

In addition, simulated data were produced using DW-POSSUM [1] (an extension of FSL POSSUM [6]), a realistic Spin-Echo EPI simulator. It takes as input tissue segmentations, MR parameters associated with these tissues and a pulse sequence, and produces an EPI image by solving the Bloch equations. We obtained the tissue segmentations by processing the anatomical scans with FSL FAST, producing probability maps for grey matter, white matter and cerebrospinal fluid. We then applied the distortion estimated by TOPUP on the real EPI data of each subject to their corresponding simulated undistorted images to create new synthetic pairs of AP and PA distorted images.

Acquired, TOPUP processed and simulated data have been used to produce two datasets, denoted as real and simulated, in order to cover various experimental configurations. A diagrammatic representation of the processing paths followed to obtain the two datasets is presented in Fig 2.

– In the real dataset: the inputs of the model are the acquired AP and PA images, the ground truth distortion fields used at training and for evaluation are the ones from TOPUP, the ground truth images used only for evaluation are the undistorted images from TOPUP. The advantage of this dataset is that it is made of real data and all the artifacts it implies. The drawback is that TOPUP is used as ground truth and therefore cannot be included in the comparison.
– In the simulated dataset: the inputs of the model are the simulated AP and PA images, the ground truth distortion fields used at training and for evaluation are the ones from TOPUP (same as for the real dataset), the ground truth images used only for evaluation are simulated non-distorted images. The advantage of this dataset is the synthetic, absolute nature of the ground truths allowing any algorithm comparison. The drawback is that the simulation process is not able to reproduce all the artifacts induced by a real acquisition.

**Fig. 2.**
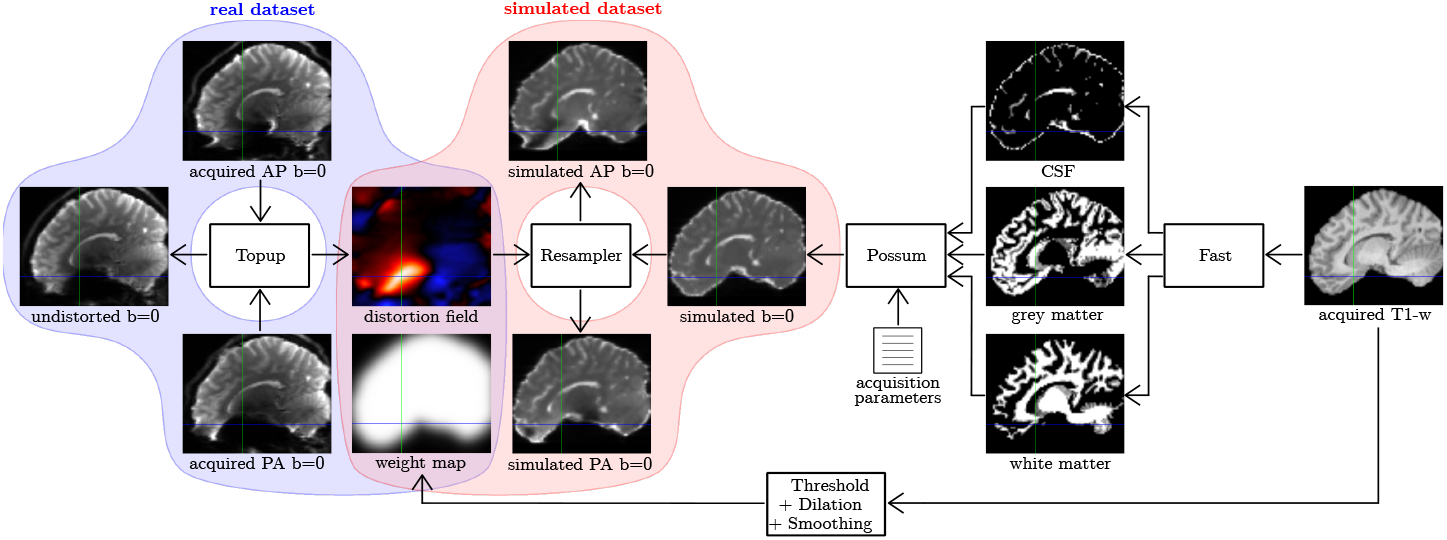
Preprocessing steps involved in creating the real and the simulated datasets.

For both, the weight maps have been computed by thresholding (binary brain mask), then dilating (3 voxels), then smoothing (Gaussian, *σ* = 6 mm) the T1-weighted images.

We divided the same way the two datasets into training (n=40), validation (n=10) and testing (n=10) samples. Although it was shown in [12] that Voxelmorph-like architectures can achieve decent registration with only ten or so training subjects.

### 4.2 Models

We compared 3 deep learning approaches: unsupervised, transformation supervised and semi-supervised. Each model is trained and assessed separately for real and simulated experiments. To form the overall loss, the unsupervised loss was attributed a weight 200 000 (unsupervised and semi-supervised models only), the supervised loss was attributed a weight 300 (supervised and semi-supervised models only) and the regularization loss was attributed a weight 1 (all models). Each model was trained for 1000 epochs, with the epoch giving the best validation loss kept. Learning rate was set to 10^-4^. The training time for each model was about 3 hours on a GPU.

### 4.3 Assessment metrics

For both real and simulated data experiments, their respective unseen test data were used to quantitatively assess the performance of the proposed semisupervised approach against the current unsupervised approach. The assessment made use of the following metrics: 1) Image fidelity: MSE between the estimated undistorted versions of the AP and PA images and the corresponding ground truth undistorted image. 2) Field fidelity: MSE between the estimated distortion field and the corresponding ground truth distortion field (expressed in fraction of the voxel size which is 2 mm). These mean measures were weighted by the weight maps to ignore the background.

## 5 Results and Discussion

Table 1 and Fig. 3 summarise the evaluation results for both real and simulated data experiments.

– The real dataset experiment uses real acquired EPI images and evaluate how close to TOPUP (ground truth) the different models behave. In terms of field fidelity, supervised and semi-supervised models show a mean voxelique MSE around 0.2 (equivalent to 0.4 mm^2^), way better than the unsupervised approach that shows an error about 4 times bigger with also a much higher variance. In terms of image similarity, the unsupervised and the semi-supervised approach show an error equivalent to 2/3 of the one of the supervised approach with akin variance. The semi-supervised approach performs well for both metrics whereas the other models present weaker results in one situation. An example case subject from the testing sample, corrected with the semi-supervised approach, is shown in Fig. 4.
– The synthetic dataset experiment uses simulated EPI images and evaluate how each model and TOPUP are able to retrieve a synthetic ground truth. It quite follow the same trend as above for the deep-learning models. TOPUP, the supervised and the supervised models show similar field fidelity that is better than the one of the unsupervised model. TOPUP, the unsupervised and the supervised models show similar image fidelity that is better than the one of the supervised model. TOPUP and the semi-supervised model performs similarly well for both metrics whereas the supervised and the unsupervised models are weaker in one situation.

**Fig. 3.**
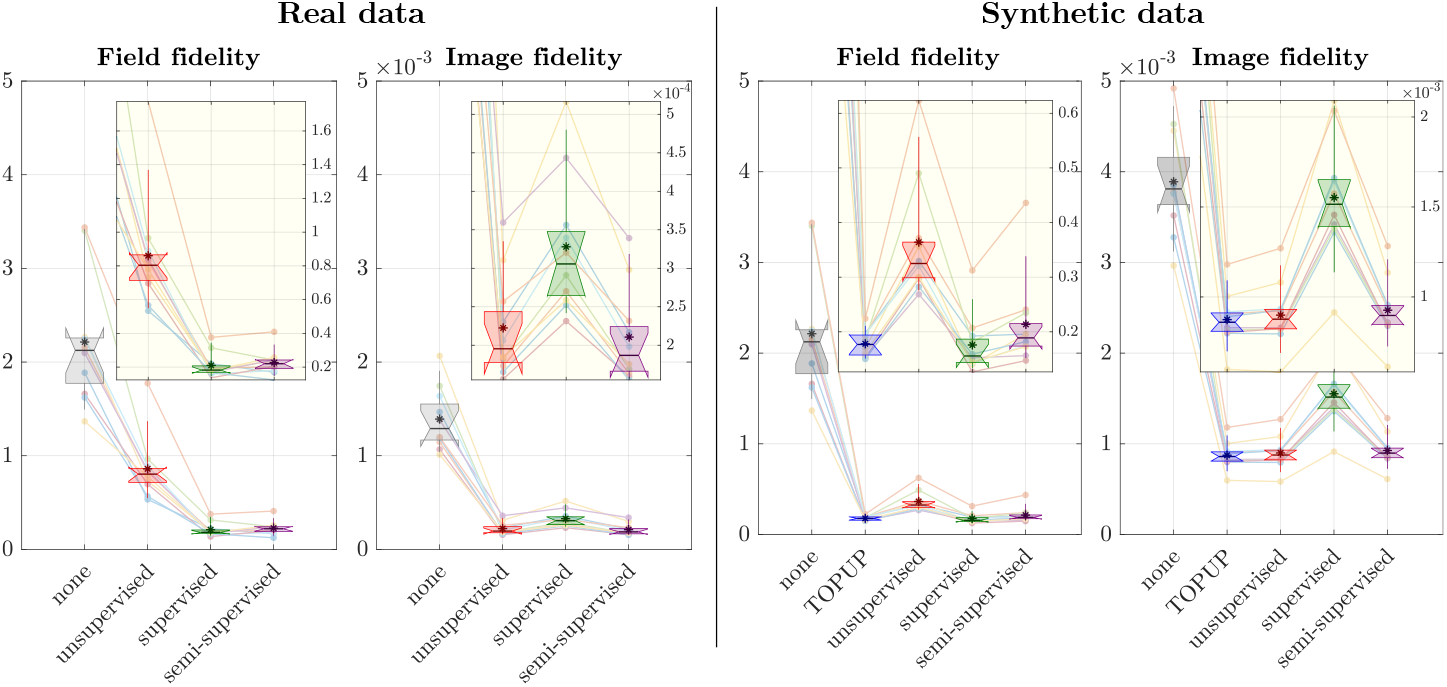
Fidelity (MSE) to ground truth distortion fields and images for the different models for real data with TOPUP as ground truth and for synthetic data. The field fidelity is expressed as a fraction of the voxel dimension (2 mm). Large frames share the same scale for global overview between experiments whereas yellowish ones are zoomed in for intra-metric, between models comparison.

**Fig. 4.**
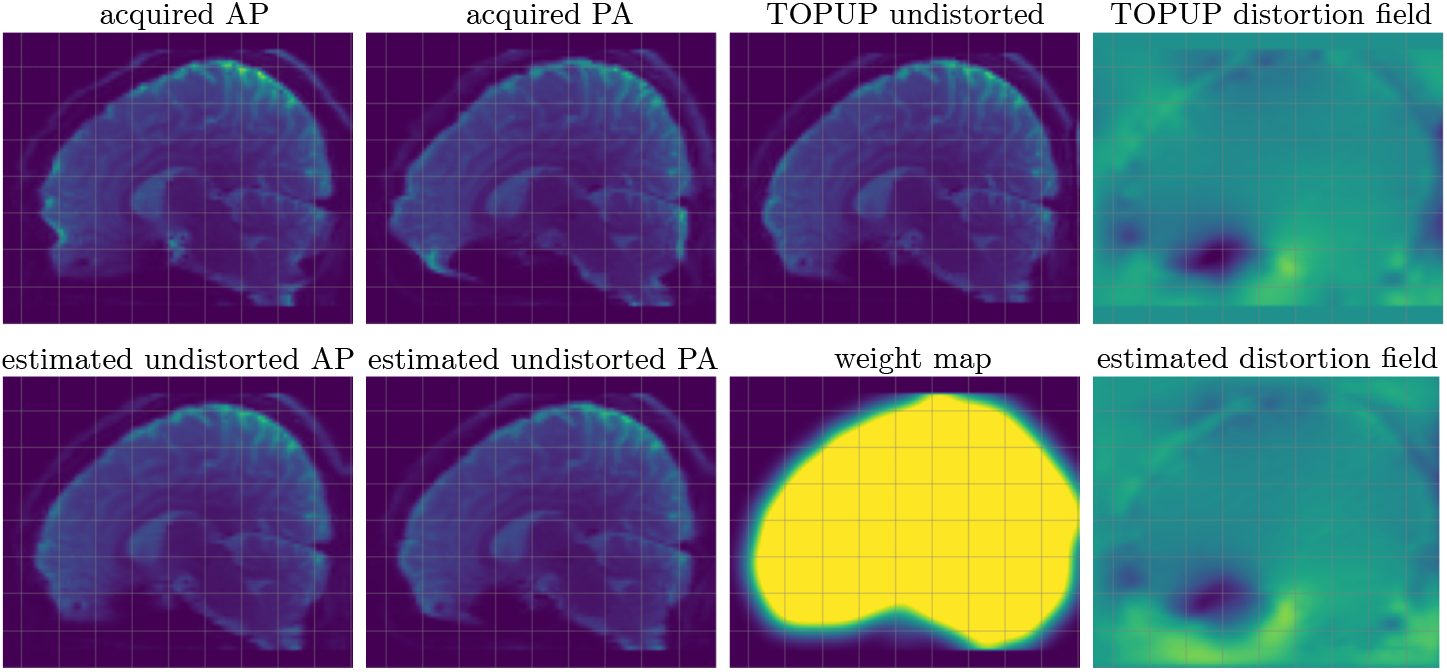
Example case from the testing sample (unseen) of the real dataset, corrected using the semi-supervised model.

**Table 1.** Fidelity (MSE: mean (std)) to ground truth distortion fields and images for the different models for real data with TOPUP as ground truth and for synthetic data. The field fidelity is expressed as a fraction of the voxel dimension (2 mm).

|  |  | none | TOPUP | unsupervised | supervised | semi-supervised |
| --- | --- | --- | --- | --- | --- | --- |
| Real dataset | Field fidelity | 2.21<br>(0.70) | ∅ | 0.86<br>(0.34) | 0.21<br>(0.07) | 0.22<br>(0.07) |
| | Image fidelity | $1.39 \cdot 10^3$<br>($3.39 \cdot 10^4$) | ∅ | $2.22 \cdot 10^4$<br>($6.76 \cdot 10^5$) | $3.27 \cdot 10^4$<br>($9.09 \cdot 10^5$) | $2.10 \cdot 10^4$<br>($6.29 \cdot 10^5$) |
| Synthetic dataset | Field fidelity | 2.21<br>(0.70) | 0.17<br>(0.02) | 0.36<br>(0.11) | 0.17<br>(0.05) | 0.21<br>(0.08) |
| | Image fidelity | $3.89 \cdot 10^3$<br>($5.94 \cdot 10^4$) | $8.73 \cdot 10^4$<br>($1.51 \cdot 10^4$) | $8.98 \cdot 10^4$<br>($1.82 \cdot 10^4$) | $1.55 \cdot 10^3$<br>($3.41 \cdot 10^4$) | $9.25 \cdot 10^4$<br>($1.81 \cdot 10^4$) |

This study show that, with proper distortion model, one can use deeplearning registration trained on some tens of subjects to rapidly correct a larger set for susceptibility-induced distortion, with results as good as TOPUP. One can use a processed subset of a dataset to include transformation supervision and improve the transformation fidelity which is a direct measure of the quality of the registration. We could have expected the semi-supervised model to perform half-way between its unsupervised and supervised counterparts in terms of field and image fidelity; it actually offers the best of both worlds.

## 6 Conclusion

We presented a semi-supervised approach for the distortion correction of EPI images with opposite PED using DL-based image registration. We compared this model with an unsupervised and a supervised one, as well as a traditional algorithm: TOPUP. By leveraging ground truth distortion transformations during training, the proposed method can produce more accurate estimate of distortion fields (direct quantitative metric) compared to unsupervised approaches at testing. It also outperforms the supervised approach for the image metric. On synthetic data, the results were similar to TOPUP but with much faster computation. The proposed model can typically be trained on a processed subsample of a dataset with an external tool to then produce distortion correction very efficiently to the rest.

## References

1. Graham M. S., Drobnjak I., Jenkinson M. and Zhang H., Quantitative assessment of the susceptibility artefact and its interaction with motion in diffusion MRI. PloS one, 12(10) (2017).

2. Gu X., Eklund A., Evaluation of Six Phase Encoding Based Susceptibility Distortion Correction Methods for Diffusion MRI. Frontiers in Neuroinformatics: vol. 13 (2019)

3. Andersson J.L., Skare S., Ashburner J., How to correct susceptibility distortions in spin-echo echo-planar images: application to diffusion tensor imaging. NeuroImage, 20(2), 870–888 (2003).

4. Duong S.T.M., Phung S.L., Bouzerdoum A., Schira M.M., An unsupervised deep learning technique for susceptibility artifact correction in reversed phase-encoding EPI images, Magnetic Resonance Imaging: Volume 71, 1–10 (2020).

5. Rohlfing T., Image Similarity and Tissue Overlaps as Surrogates for Image Registration Accuracy: Widely Used but Unreliable, IEEE Transactions on Medical Imaging, vol. 31, no. 2, pp. 153–163, (2012)

6. Drobnjak I., Gavaghan D., Süli E., Pitt-Francis J., Jenkinson M., Development of a functional magnetic resonance imaging simulator for modeling realistic rigid-body motion artifacts. Magnetic resonance in medicine, 56(2), 364–380 (2006).

7. Jezzard P. and Balaban RS., Correction for geometric distortion in echo planar images from B0 field variations. Magnetic resonance in medicine: vol. 34, 1: 65–73 (1995).

8. Morgan PS, Bowtell RW, McIntyre DJ, Worthington BS., Correction of spatial distortion in EPI due to inhomogeneous static magnetic fields using the reversed gradient method. Journal of magnetic resonance imaging: vol. 19, 4 (2004): 499–507.

9. Chang H., and Fitzpatrick JM., A technique for accurate magnetic resonance imaging in the presence of field inhomogeneities. IEEE transactions on medical imaging: vol. 11, 3 (1992): 319–29.

10. Fu Y., Lei Y., Wang T., Curran W.J., Liu T., Yang X., Deep learning in medical image registration: a review. hys Med Biol, (2020).

11. Haskins G., Kruger U., Yan P., Deep learning in medical image registration: a survey. Machine Vision and Applications 31, 8 (2020).

12. Balakrishnan G., Zhao A., Sabuncu M.R., Guttag J.V. and Dalca, A.V., Voxel-Morph: A Learning Framework for Deformable Medical Image Registration. IEEE Transactions on Medical Imaging, 38, 1788–1800 (2019).

13. Dalca A.V., Balakrishnan G., Guttag J.V. and Sabuncu, M.R., Unsupervised Learning of Probabilistic Diffeomorphic Registration for Images and Surfaces. Medical image analysis, 57, 226–236 (2019).

14. Ronneberger O., Fischer P., Brox T., U-Net: Convolutional Networks for Biomedical Image Segmentation. Medical Image Computing and Computer-Assisted Intervention – MICCAI (2015). LNCS, vol 9351.

